# Integrins control tissue morphogenesis and homeostasis by sustaining the different types of intracellular actin networks

**DOI:** 10.1101/639427

**Authors:** Carmen Santa-Cruz Mateos, Andrea Valencia-Expósito, David G. Míguez, Isabel M. Palacios, María D. Martín-Bermudo

**Author notes:** Communicating author: M. D. Martín-Bermudo.

## Abstract

Forces generated by the actomyosin cytoskeleton are key contributors to the generation of tissue shape. Within the cell, the actomyosin cytoskeleton organizes in different types of networks, each of them performing distinct roles. In addition, although they normally localize to precise regions of the cells, they are rarely independent and often their dynamics influence each other. In fact, the reorganization of a given structure can promote the formation of another, conversions that govern many morphogenetic processes. In addition, maintenance of a specific actomyosin network organization in a differentiated tissue might be equally important. Failure to do so could lead to undesired cell state transitions, which in turn would have drastic consequences on the homeostasis of the tissue. Still, little is known about the mechanisms that ensure controlled transitions between actomyosin networks during morphogenesis or their maintenance in a differentiated tissue. Here, we use the *Drosophila* follicular epithelium to show that cell-ECM interactions mediated by integrins are necessary for the establishment and maintenance of the different actomyosin networks present in these epithelial cells. Elimination of integrins in a group of follicle cells results in changes in the F-actin levels and physical properties of their intracellular actomyosin networks. Integrin mutant follicle cells have reduced number of basal stress fibers. They also show increased cortical F-actin levels and tension, which interferes with proper basal surface growth. Finally, clonal elimination of integrins also triggers non-autonomous behavioural changes in neighbouring wild types cells, which now reorganize their actin cytoskeleton and spread and overlay the mutant ones. Based on these results, we propose that cell-ECM interactions mediated by integrins regulate epithelia morphogenesis and homesostasis by preserving the different types of intracellular actin networks.

## Introduction

Forces generated by F-actin networks are important contributors to the generation of cell and tissue shape. The architecture and mechanical properties of the F-actin network are modulated by myosin II motors and actin binding proteins (reviewed in ^1^. The molecular composition of contractile actin networks and bundles is highly conserved among eukaryotic species ^2^. Nevertheless, their organization and dynamics change across different cell types, their position within the cell and the differentiation state of the cell.

There are two main ways in which actomyosin networks can be organized within the cell, as cortical two-dimensional meshworks below the plasma membrane or as stress-fibers. Studies over the last decade have assigned distinct roles for these two types of networks. Thus, while pulsatile contraction of cortical actomyosin meshworks has been mainly implicated in the cell shape changes underlying key morphogenetic processes, such as gastrulation and neural tube formation (reviewed in ^3^, stress fibers have been largely involved in cell adhesion, migration and mechanosensing ^4^. During morphogenesis, cells change the way they organize their actin networks in response to intracellular signals. For example, a change in actin organization from a cortical rearrangement into stress fibers is observed when cells exit mitosis or naïve pluripotency or during epithelial to mesenchyme transition (reviewed in ^5^. In addition, transitions in actin organization can also depend on the way cells interact with each other or with the extracellular environment. Thus, while cell-cell interactions promote cortical actin organization, cell-ECM adhesion stimulates stress fibers formation. During morphogenesis, transitions between these two different actin networks need to be finely regulated in space and time, as misplaced or untimely transitions could affect the proper formation of organs and tissues. Finally, maintenance of a specific actomyosin network organization in a differentiated cell is crucial for proper organ function. Failure to do so could trigger changes in the cell stage, which could ultimately have drastic consequences on the homeostasis of a tissue. Still, little is known about the mechanisms that ensure controlled transitions in actin reorganization during morphogenesis or maintenance of a specific actin network in a differentiated cell.

The follicular epithelium of the adult *Drosophila* ovary provides an excellent system to study the contribution of cell-ECM interactions to actin network organization during morphogenesis. The *Drosophila* ovary is composed of 16-18 tube-producing eggs called ovarioles ^6^. Each ovariole contains a germarium at their anterior end and progressively older egg chambers at their posterior end. Each egg chamber is composed by 15 nurse cells and one oocyte enveloped by a single layer of follicle cells (FCs), termed the follicular epithelium (FE) ^7^. Newly formed egg chambers are round and go through 14 developmental stages (from S1 to S14) to eventually give rise to mature eggs, which are elongated anterior-posteriorly ^7^. At the time that the egg chamber buds from the germarium, approximately 80 FCs enclose the germline cyst ^7^. FCs continue to divide mitotically until S6 when they exit the mitotic cycle and switch to an endocycle ^8^. Between S7-10, FCs undergo three rounds of endoreplication, become polyploid and increase their size. The apical side of FCs faces the germline, while their basal surface contacts a specialized ECM called basement membrane, which encapsulates the egg chamber ^9^. On the basal surface of FCs, F-actin organizes not only on the two most common types of networks, a cortical meshwork and polarized stress fibers ^10^, but also in planar polarized whip-like structures that emerge predominantly at tricellular junctions ^11^, each of them with distinct roles (see below). Stress fibers change their morphology and orientation throughout oogenesis ^10^. In addition, they terminate at sites containing integrins and components of focal adhesion, such as paxillin, talin, PINCH and ILK (Fig.1A, B and ^10, 12, 13^. Furthermore, integrins are required for stress fibers attachment and control of F-actin levels. FCs lacking integrins display a 33% increase in F-actin levels, which have led to propose that integrins are required to reduce fibers numbers ^10^. This differs from the proposed role for integrins in formation of stress fibers in cells in culture ^14^. In contrast, but in agreement with results from cell culture experiments, basal myosin levels were shown to decrease in FCs with reduced levels of integrins or its interacting protein Talin ^13, 15^. Thus, the role of integrins on the organization of actomyosin fibers on the basal side of FCs remains controversial. The actomyosin fibers present on the basal side of FCs undergo oscillating contractions ^13, 16^. Furthermore, these contractions and their coupling to the ECM are required for the cell shape changes underlying egg elongation ^12, 13, 17, 18^. However, how basal actomyosin fibers and their attachment to the ECM control cell shape changes remains poorly understood. In summary, F-actin organizes in three different types of networks at the basal side of FCs. The mechanisms regulating the assembly and maintenance of these different networks and their roles during FE morphogenesis and homeostasis remain unclear.

**Figure 1.**
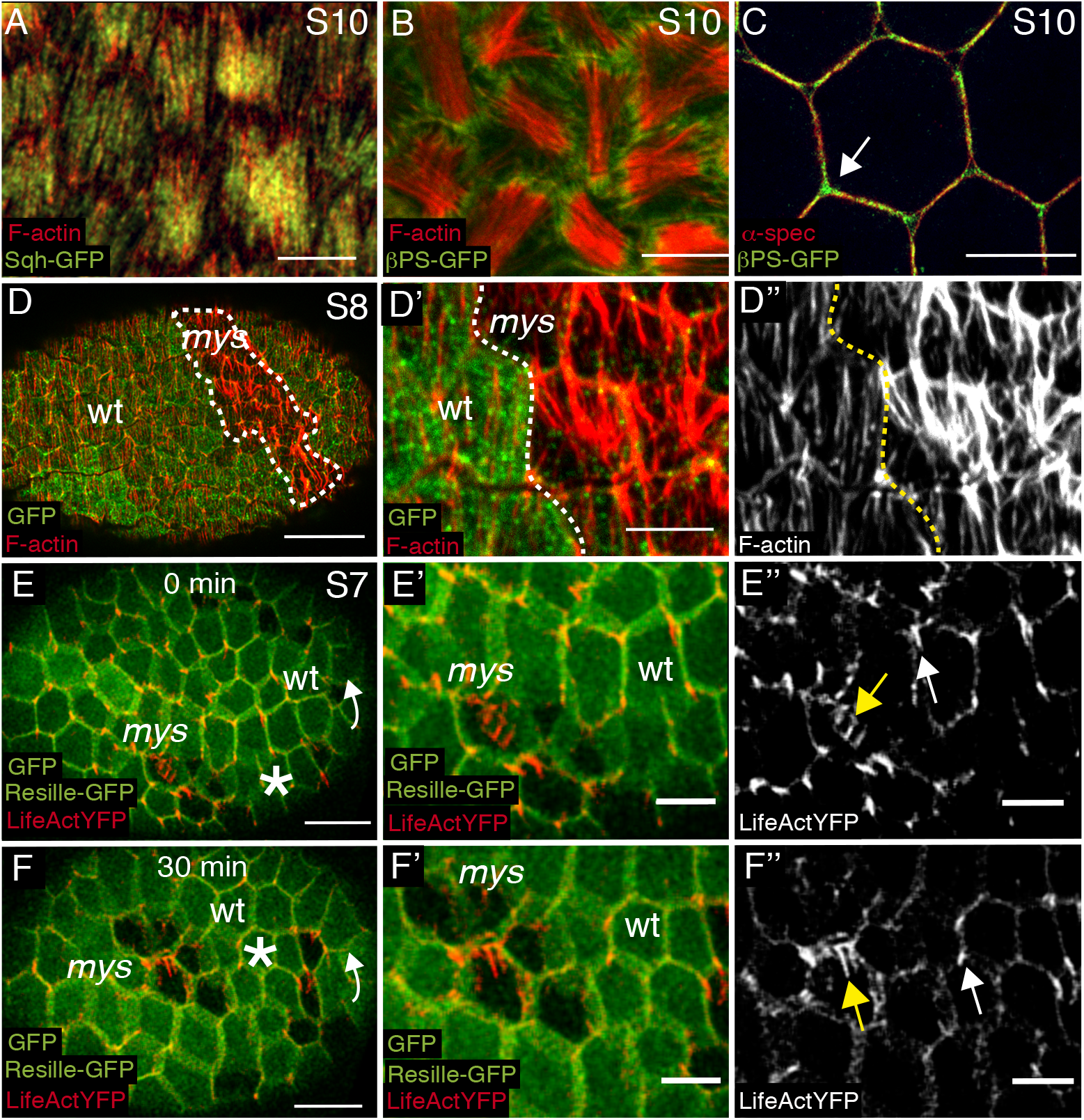
Elimination of integrins from FCs increases actin-rich whip-like structures content. **(A, B)** Basal surface views of S10 FCs expressing (A) Sqh-GFP (green) or (B) βPS-GFP (green) to visualize myosin or integrins, respectively, stained with Rhodamine-Phalloidin (red) to detect F-actin. **(B)** Integrins localize at actin stress fibers tips. **(C)** Sagittal plane of S10 FCs expressing βPS-GFP (green) stained with anti-GFP and an antibody against the cortical protein α-spectrin (red). Integrins are found cortically and at tricellular junctions (arrow). **(D)** Basal surface view of **a** mosaic S8 egg chamber containing *mys* mutant FCs and stained for anti-GFP (green) and Rhodamine Phalloidin (red). *mys* mutant FCs (labelled by de absence of GFP, GFP^-^) show higher levels of basal F-actin than controls (GFP^+^). **(D’, D’’)** Magnifications of the area within the dashed line in **D. (E, F)** Confocal images taken with 30 minutes difference of a live S7 mosaic egg chamber expressing lifeactin-YFP (red) and carrying *mys* mutant clones. *mys* FCs (GFP^-^, yellow arrow) show an increase in actin-rich whip-like structures (arrowhead) compared with controls (GFP^+^, white arrow). Arrows in **E** and **F** indicate direction of egg chamber rotation. Asterisks label a cell as a reference for the rotation. Scale bar, 10μm. **(E’, E’’, F, F’’)** Magnifications of **E** and **F** respectively. Scale bars, 10μm (**A-C**), 20μm (d) and 5μm (**D’-F’’**).

Here, we show that integrins are necessary for the establishment and maintenance of the three different actin networks present in the FCs. Elimination of integrins results in changes in F-actin levels and physical properties of these meshworks. This leads to severe cell-autonomous and non-autonomous consequences for this epithelium. On one hand, the basal surface of integrin mutant follicle cells does not reach its normal size. On the other hand, elimination of integrins in a limited group of cells induces a behavioural change in neighbouring wild types cells, which now change their actin cytoskeletal organization and spread and overlay the mutant ones. Based on these results, we propose that cell-ECM interactions mediated by integrins regulate epithelia morphogenesis and homeostasis by sustaining the different types of intracellular actin network.

## Results

### Elimination of integrins from FCs increases actin-rich whip-like structures

Integrins are heterodimeric receptors composed of an α and a β subunit. While in vertebrates there are at least 8 β subunits and 18 α subunits, in *Drosophila* there are only two β subunits, βPS and βν, and five α subunits, αPS1 to αPS5. We have previously shown that elimination of *myospheroid* (*mys*), the gene encoding the βPS subunit, from FCs removes all integrins present in the ovary and results in multilayering, polarity and differentiation defects at both poles of the egg chamber ^19–21^. Previous studies have shown that integrins and associated proteins localize at actin stress-fiber ends ^10, 12^, Fig.1A, B). In addition, we found that integrins also accumulated cortically, localizing with proteins associated to cortical F-actin, such as spectrins (Fig. 1C, ^22^. Finally, integrins were also found highly enriched at tricellular junctions (arrowhead in fig.1C).

Next, we analyzed basal actomyosin organization in FCs homozygous mutant for the null allele *mys*^XG43 23^. In agreement with previous results ^10^, we found that FCs lacking βPS showed increased levels of F-actin at S8 (Fig.1D-D’’). However, the increase in F-actin observed in *mys*^XG43^ FCs seemed associated with the whip-like structures found at tricellular junctions rather than with the stress fibers ^11^. We tested this by analyzing actin dynamics in control and mutant FCs *in vivo*. To do this, we generated transgenic flies expressing LifeActin-YFP under a ubiquitous promoter, Ubi-LifeActinYFP (see Materials and Methods for details), which marks cortical F-actin and all actin-rich protrusions, including the whip-like structures (Movie S1). Indeed, we found an increase in whip-like structures in S7 *mys*^XG43^ FCs compared to controls (Movie S1, Fig.1E, F). The ectopic whip-like structures found in *mys*^XG43^ FCs showed a dynamic behavior similar to that observed in neighboring control FCs, i.e. propel contrary to the direction of egg chamber rotation (Movie 1). However, and in contrast to what happens in control cells, the whip-like structures did confine to tricellular junctions in mutant FCs. Basal whip-like structures are enriched in F-actin and do not contain myosin ^11^. Analysis of myosin distribution and dynamics in wild type and *mys*^XG43^ FCs *in vivo*, using the green fluorescent protein GFP fused to the myosin regulatory light chain Spaghetti-Squash (Sqh-GFP, ^24^, showed no increase in basal myosin levels in the integrin mutant FCs (Fig.S1 and Movie S2). These results support our hypothesis that the rise in basal F-actin levels observed in integrin depleted FCs results in an increase in the whip-like structures and not to a rise in the number of actomyosin stress fibers.

Next, we decided to analyze in more detail stress fiber formation, organization and maintenance in the absence of integrins.

### Integrins are required for proper actomyosin fiber formation, maintenance and dynamics

Stress fibers and focal adhesions present on the basal side of FCs are analogous to those formed by many different cell systems in culture, such as fibroblasts, epithelial and endothelial cells ^4^. Studies using these cell culture systems have highlighted the existence of a mechanical crosstalk and interdependence between the physically coupled stress fibers and focal adhesions. Stress fibers diminish when their focal adhesions disassemble during cell migration and vice versa, focal adhesions disassemble when stress fibers are disrupted (reviewed in ^25^.

Although present from the beginning, stress fibers on the basal side of FCs change their organization and orientation throughout oogenesis ^10^. At S6 new F-actin filaments become visible. From S8, these filaments extend until S9, when they cover the basal surface of the cells. Up to S10A, fibers orient perpendicular to the long axis of the egg chamber. By S10B they become thicker and less well oriented ^10^. As mentioned in the introduction, the role of integrins on the morphogenesis of the actomyosin fibers present on the basal side of FCs remains somehow contentious ^10, 13, 15^. Thus, we decided to study this in more detail by analysing the density and morphology of the actomyosin fibers throughout FC development, using Rhodamine-phalloidin and Sqh-GFP to visualize F-actin and myosin, respectively (Fig.2, Fig.S2). We found that already at S8 the density of actomyosin fibers was reduced in *mys*^XG43^ FCs compared to controls (Fig.2A, A’, Fig.S2A, A’). This phenotype worsened as oogenesis progressed (Fig.2B, B’, Fig.S2B, B’), so that by S10 basal stress fibers were hardly detectable (Fig.2C, C’ and Fig.S2C, C’). To better visualize, them we used SiR-Actin and Stimulated Emission Depletion (STED) microscopy, which allows the generation of super-resolution images of F-actin networks ^26^. This showed that indeed the stress fibers of S10 mutant FCs were further reduced in number and length compared to controls (Fig.2C, C’). We next decided to quantify this phenotype by measuring the number and morphology of stress fibers at S9, when they are fully extended in wild type FCs and still visible in the *mys*^XG43^ mutant FCs. To do this, we measured F-actin staining intensity across a 2μm bar centered at the basal side of FCs and identified peaks in which the fluorescence intensity exceeded one standard deviation below the mean intensity in the wild type (see Material and Methods). We found that the number of peaks per micrometer was lower in *mys*^XG43^ FCs compared to controls (Fig.2D). Likewise, the overall intensity of F-actin in stress fibers in the mutant FCs was reduced by 60% with respect to controls (Fig.2E). Finally, using live imaging of mosaic S10 egg chambers expressing either our Ubi-LifeActinYFP (Fig.S3, Movie 3) or Sqh-mCherry (Fig.S4, Movie 4), the membrane marker Resille-GFP (^27^ and containing *mys*^XG43^ FCs, we found that, in agreement with previous results ^15^, the preferential pulsation period of both F-actin and myosin in mutant cells was reduced compared with control cells, 4.5min versus 6.6 (Fig.S3C-E) and 4.05min. versus 6.90min (Fig.S4C-E), respectively (Movies 3 and 4). In addition, we found that mutant cells displayed a more stochastic behavior than controls (Fig.S3C and Fig.S4C). Finally, we challenged our previous model and tested if reduced F-actin levels could lead to shorten and more stochastic oscillations, as well as reduced myosin levels ^16^. We found this was the case (Fig.S3F-H), proving the robustness of the model. Altogether, our results show that integrins regulate the formation, maintenance and dynamics of the basal actomyosin network.

**Figure 2.**
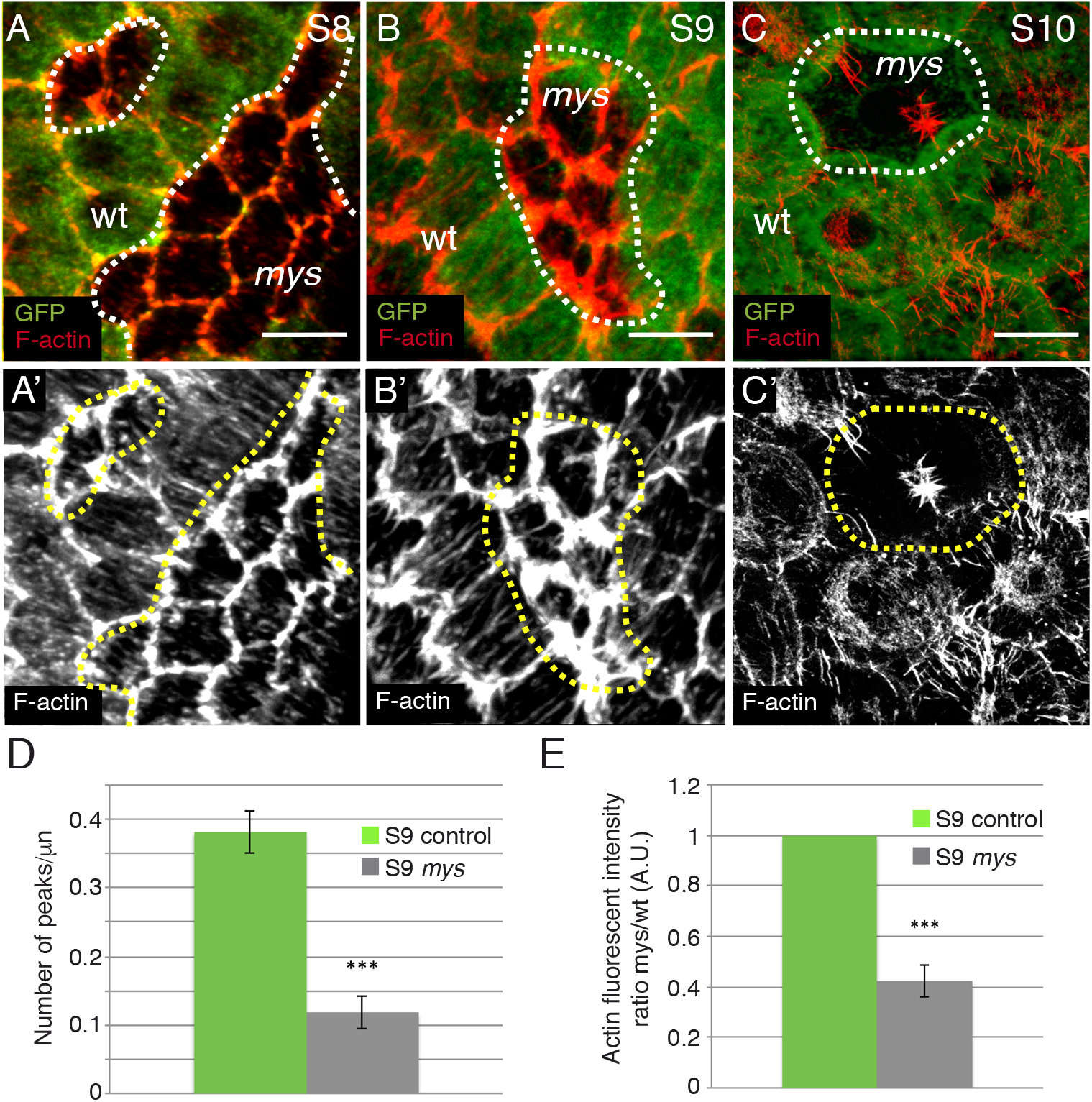
Integrins are required for proper stress fiber formation and maintenance. **(A, B, C)** Basal surface of mosaic S8 (**A**), S9 (**B**) and S10 (**C**) egg chambers stained with anti-GFP (green) and Rhodamine Phalloidin (red) to visualize F-actin. (**A, A’**) S8 *mys* mutant FCs show reduced stress fibers number compared with controls. (**B, B’**) At S9 stress fibers are hardly visible in *mys* mutant FCs (GFP^-^). **(C, C’)** By S10, only a few short actin fibers are observed in the mutant cells. **(D)** Quantification of the number of actin fibers per μm in control and *mys* mutant FCs. **(E)** Quantification of the ratio of total fluorescence intensity of F-actin present in stress fibers in control and mutant FCs. The statistical significance of differences was assessed with a t-test, *** P value < 0.0001. All errors bars indicate s. e. Mean of n= 25 FCs, assessed over 8 independent S9 egg chambers. Scale bars, 5 μm.

### Integrins regulate cortical F-actin levels and tension

Adherent mouse embryonic fibroblasts that gradually detach from the substrate redistribute their F-actin from stress fibers to a more cortical position. Given that stress fibres were reduced in S9 *mys*^XG43^ FCs (Fig.2), we sought to determine whether cortical actin was affected in these mutant cells. We found elevated levels of cortical F-actin in the mutant FCs compared to controls (Fig.3A, A’). To quantify this, we measured cortical F-actin intensity along cell-cell contacts, using an approach similar to the one described above (see Materials and Methods). Indeed, we found a 2 fold increased in cortical F-actin in *mys*^XG43^ FCs compared to their wild type counterparts (Fig.3B, C). The increased levels of cortical F-actin found in mutant FCs was specific to the basal side, as no difference was found apically (Fig.S5A, A’). We next tested whether the defects in cortical F-actin found on the basal side of mutant FCs was cell autonomous. To do this, we generated large clones and compared cortical F-actin in mutant cells surrounded by either control or mutant cells. We found that cortical F-actin levels were increased in all mutant cells, suggesting that the defects in cortical F-actin due to elimination of integrins are cell autonomous (Fig.S5B, B’).

**Figure 3.**
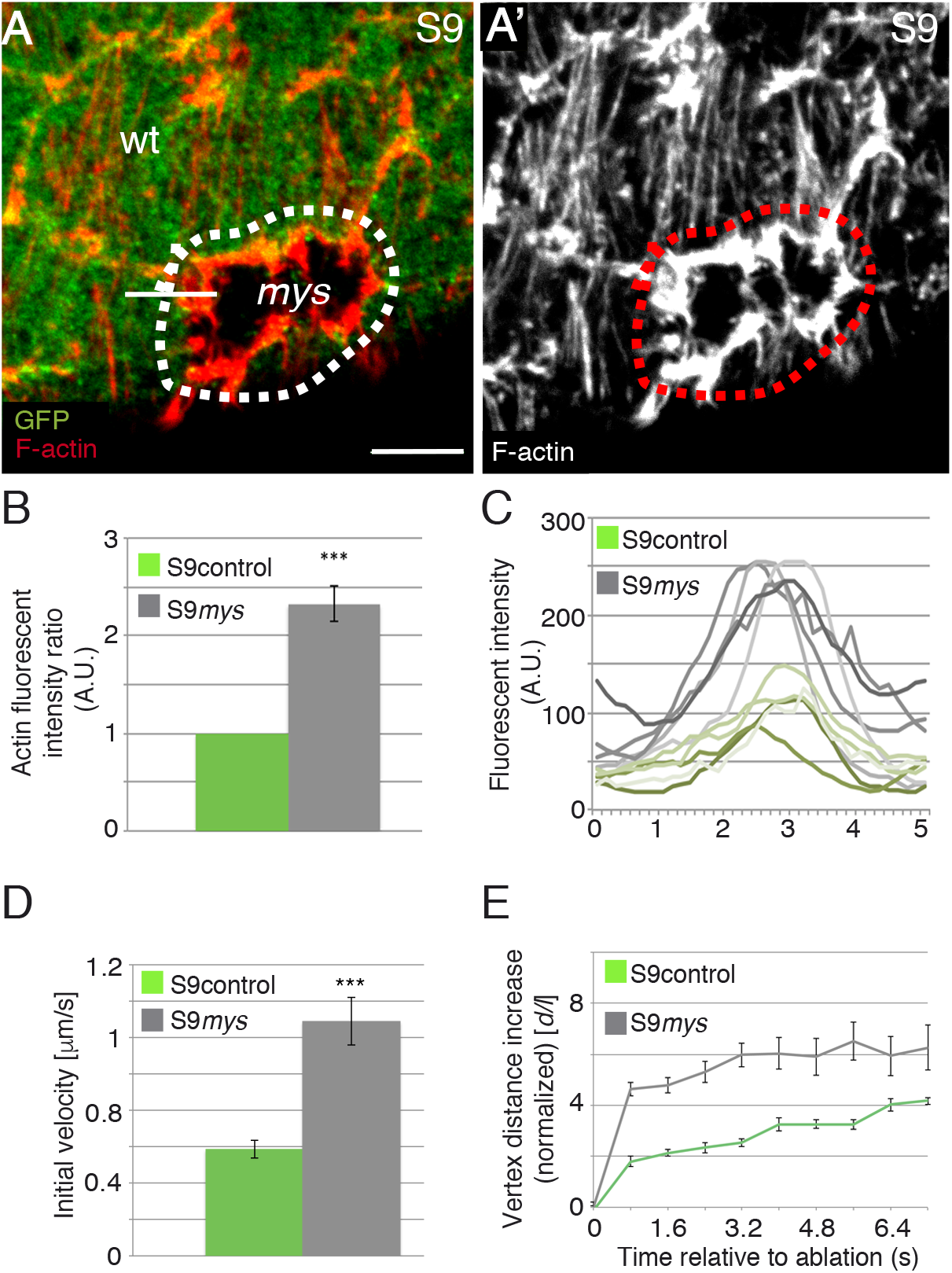
Abscence of integrins increase both cortical F-actin levels and tension. **(A, A’)** Basal surface view of a mosaic follicular epithelium stained for anti-GFP (green) and Rhodamine Phalloidin to detect F-actin (red). **(B)** Quantification of cortical actin intensity in *mys* mutant FCs (GFP^-^) with respect to controls (GFP^+^). **(C)** Histogram of relative fluorescent intensities of F-actin across the cell cortex, as indicated with a straight white line in (**A**). Five most representative curves were plotted in both control (green) and *mys* mutant (grey) FCs. Mean of n= 22 FCs from 5 different egg chambers. **(D, E)** Quantification of initial velocity of vertex displacement (**D**) and vertex displacement over time (**E**) after photoablation at the basal side of control (green) and *mys* mutant (grey) FCs. The statistical significance of differences was assessed with a t-test, *** P value < 0.0001. All errors bars indicate s. e. Quantifications were done in n= 15 FCs from 15 independent S9 egg chambers. Scale bars, 5 μm.

Laser ablation of cell-cell boundaries is as an effective tool to measure actomyosin contractility-based intracellular tensions ^28^. To explore whether the increase in cortical F-actin found on the basal side of S9 mutant FCs changed the contractility state of these FCs, we performed laser ablation experiments using a UV laser beam to sever plasma membranes and the cortical cytoskeleton on the basal side of both control and *mys*^XG43^ mutant FCs (see Materials and Methods). The behavior of cell membranes, visualized with Resille-GFP (^27^, was monitored up to ten seconds after ablation. As a consequence of the cut, cortical tension relaxed and the distance between the cell vertices at both sides of the cut increased. As velocity of retraction is affected by cytoplasmic viscosity ^29^, we assumed viscosity in *mys*^XG43^ and wild type FCs to be the same. To minimize potential effects due to the anisotropic distribution of forces in the FE, cuts were all made perpendicular to the AP axis and in the central region of fifteen independently cultured egg chambers. Furthermore, only vertices initially separated by similar distances were considered. We found that the initial velocity of vertices displacement in *mys*^XG43^ mutant FCs was two times higher than that found in control cells (Fig.3D). In addition, vertex displacement over time was also significantly greater in mutant cells compared to controls (Fig.3E). These results demonstrate that cell membranes are under higher tension in integrin mutant cells.

Altogether, these results strongly suggest that integrins regulate basal cortical F-actin levels and cortical tension in FCs.

### Morphological consequences of integrin elimination in FCs: defective basal surface expansion

Here, we show that elimination of integrins in FCs results in an increase in cortical F-actin and a reduction in the density of stress fibers specifically on the basal side of FCs ^30^. Results from cell culture experiments have led to the proposal that the interplay between cell-ECM adhesion, actomyosin contractility and cortical actin cytoskeleton define cell morphology. Thus, we next tested whether elimination of integrins from FCs affected their morphology. FCs remain cuboidal until S8. From this stage onwards, the follicular epithelium reorganizes, so that anterior terminal FCs flatten and form a squamous epithelium overlying the nurse cells, while the main body and posterior FCs adopt a columnar shape covering the oocyte ^7, 31^. We have previously shown that mosaic egg chambers containing integrin mutant FCs developed stratified epithelia at both poles ^21^. We also showed that mutant cells in ectopic layers showed polarity and cell shape defects ^20, 32^. These results have led us to propose that integrins might play a role in regulating cell shape and polarity. However, as this analysis was mainly performed in mutant cells within a multilayer, the changes in polarity and shape observed could be in part due to loss of contact with the germline. Thus, to address a direct role for integrins in FC shape, we focused our analysis on main body columnar cells. Using Resille-GFP to delineate the cortex of FCs, we found that the basal surface of the mutant FCs was smaller than that of control cells (Fig.4A). However, we did not find any difference in apical area or height between control and mutant FCs (Fig.4B, C).

**Figure 4.**
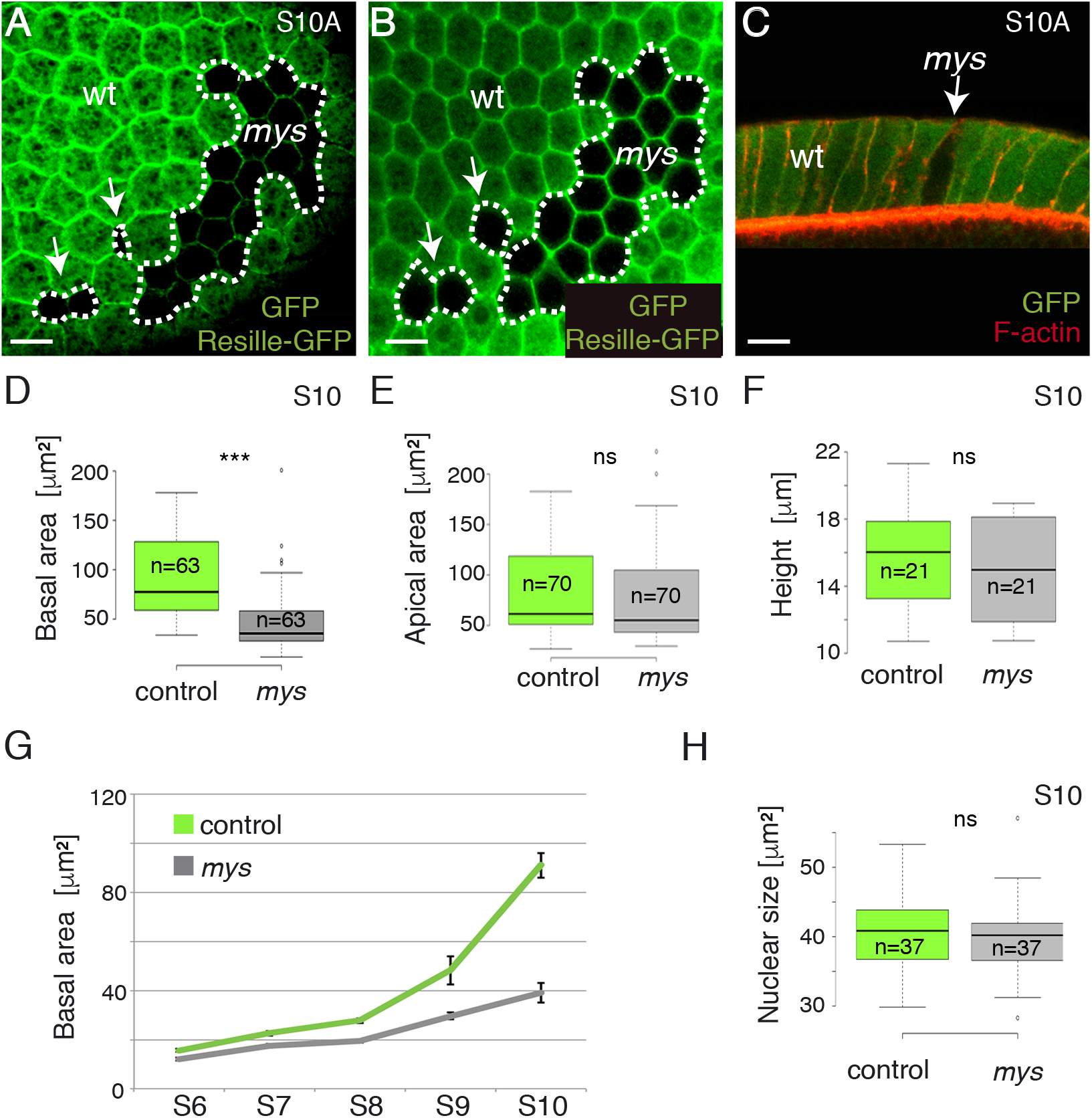
*mys* mutant FCs show defective basal surface expansion. Basal **(A)** and apical **(B)** surface views of a S10A mosaic egg chamber containing *mys* FCs and expressing the cell membrane marker Resille-GFP, stained with anti-GFP. **(C)** Lateral view of a mosaic follicular epithelium stained with anti-GFP (green) and Rhodamine Phalloidin to detect F-actin (red). **(D, E, F)** Box plots of the basal surface **(D)**, apical surface **(E)** and height **(F)** of control (green) and *mys* mutant (grey) FCs at S10. The statistical significance of differences was assessed with a t-test, *** P value < 0.0001. n is as indicated for both control and mutant FCs. **(G)** Quantification of the basal area of control (green) and *mys* mutant (grey) FCs at different stages of development. **(H)**. Box plot of the nuclear size of control (green) and *mys* mutant (grey) FCs. Error bars represent s.e. Quantifications were done in n>60 FCs, in at least 5 independent egg chambers. Scale bars, 10μm.

When cultured cells detach from the ECM, tensile loads on the cytoskeleton become unbalanced, stress fibers contract and cells shrink and adopt a rounded morphology ^33^. In this context, the reduced basal surface found in S10A *mys*^XG43^ FCs could be due to cell shrinkage. Alternatively, as FCs surface expand during stages 7 to 10 ^7, 31^, the diminished basal surface observed in the mutant FCs could also be due to defective surface expansion. To distinguish between these two possibilities, we measured surface area of control and mutant FCs throughout oogenesis. We found that the basal surface of mutant cells did not increase at the same rate as that of controls (Fig.4G). As mentioned in the introduction, between S7 and S10, FCs undergo three rounds of endoreplication and increase their size ^8^. We have previously shown that *mys*^XG43^ FCs located in ectopic layers fail to undergo proper mitotic to endocycle switch ^34^. However, we found here that the nuclear size of columnar *mys*^XG43^ FCs in contact with the germline was similar to that of controls (Fig.4H), in agreement with previous findings showing a normal BrdU incorporation pattern in mutant FCs in contact with the germline ^34^. Altogether, these results lead us to propose that the reduced basal surface observed in the mutant cells is not a failure in global growth, but a specific requirement of integrins for proper expansion of the basal surface of epithelial cells.

### Integrins control basal surface area by regulating cortical F-actin levels

Both stress fibers and cortical actin have been proposed to determine cell shape (reviewed in ^35^. Integrin mutant FCs show reduced basal surface and stress fiber number as well as increased F-actin levels in the cortex. The Abelson interacting protein, Abi, is a component of the WAVE regulatory complex that control actin dynamics. Its elimination in FCs results in the loss of cortical F-actin, with no obvious effect on the number of stress fibers ^11, 36^. To investigate whether the failure in basal area growth found in the absence of integrins was due to increase cortical F-actin, we expressed an RNAi against *abi* in integrin mutant FCs. We found that expression of *abi* RNAi in integrin FCs rescues both the increased levels of cortical F-actin and the reduced basal surface (Fig.5). As expected, the density and number of stress fibers was not changed. This result suggests that the role of integrins in controlling the basal surface area of FCs depends on their role as regulators of cortical F-actin levels and tension. To further support this idea, we interfered with actomyosin contractility by expressing a dominant negative form of *zipper* (*zip^DN^*), the gene encoding for the heavy chain of the non-muscle myosin in *Drosophila*, in integrin mutant FCs. Control FCs expressing *zip^DN^* displayed decreased F-actin levels both in the cortex and in stress fibers (Sup. Fig.6). Expression of *zip^DN^* in integrin mutant FCs restored cortical F-actin levels and basal surface area, even though it reduced further stress fibers number (Sup. Fig.6).

**Figure 5.**
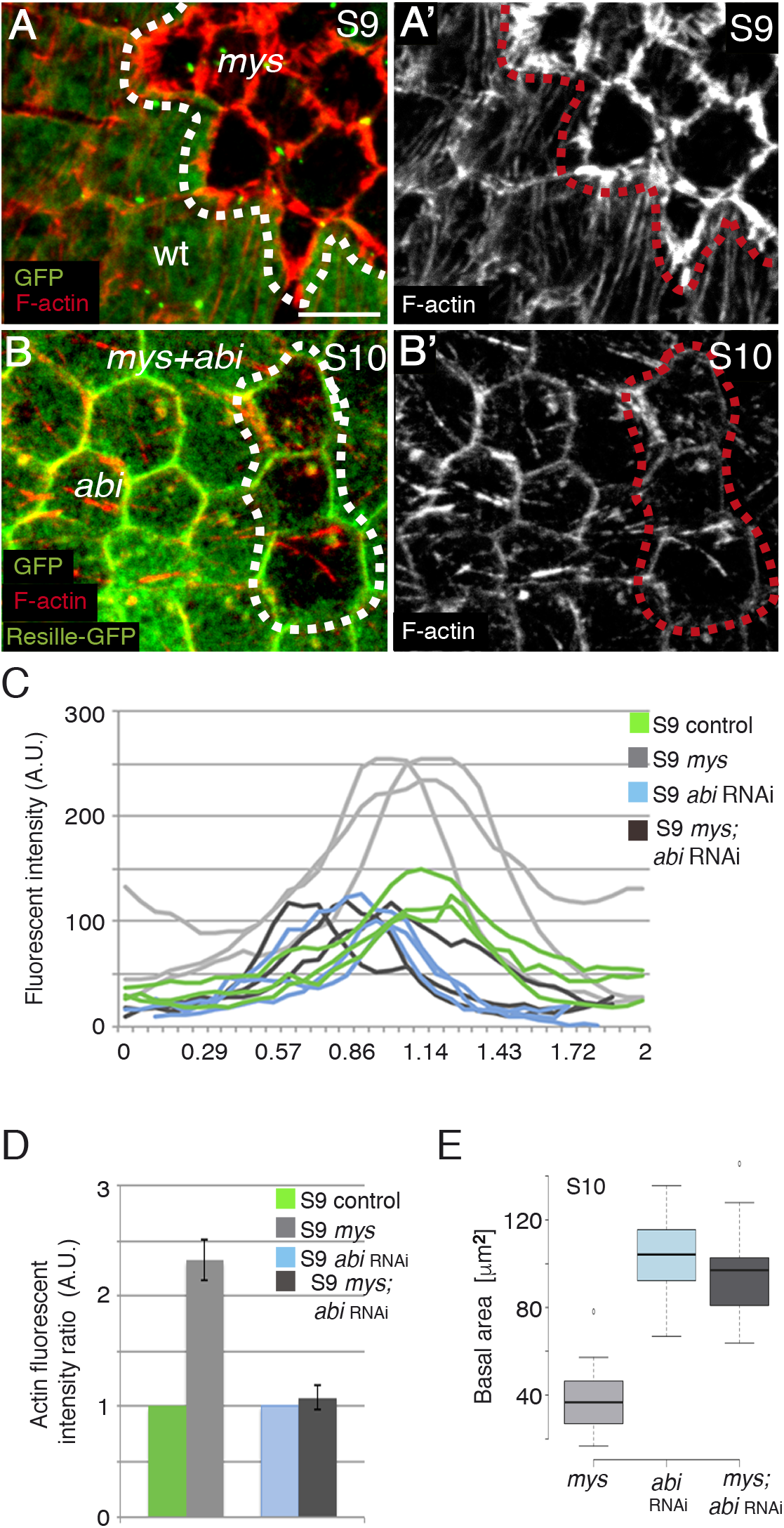
Integrins regulate cell surface growth by controlling cortical F-actin levels. **(A, A’, B, B’)** Basal surface view of a mosaic follicular epithelium containing *mys* FCs **(A, A’)** and *mys* mutant FCs expressing an *abi* RNAi **(B, B’)**, stained for anti-GFP (green) and Rhodamine Phalloidin to detect F-actin (red). **(C)** Histogram of relative fluorescent intensities of F-actin across the cell cortex of control (GFP^+^ in **A**), *mys* (GFP^-^ in **A**), *abi* RNAi (GFP^+^ in **B**) and *mys*+*abi* RNAi (GFP^-^ in **B**) FCs. Three most representative curves were plotted in all cases. **(D)** Quantification of cortical actin intensity of the indicated genotypes. **(E)** Quantification of the basal area of the indicated genotypes at S10. Mean of n= 22 FCs from 5 different egg chambers.

Finally, the function of the WAVE regulatory complex is also required for the formation of the junctional whip-like protrusions in FCs. In fact, suppression of the function of the central WRC component Abi (or Fat) results in the elimination of these structures. We found that reduction of Abi expression in *mys*^XG43^ FCs also resulted in loss of the actin-rich structures (Fig.S7).

### Integrin mutant FCs induce a non-cell autonomous cytoskeletal reorganization and behavior in neighboring wild type cells

When examining the growth of the basal surface of FCs throughout oogenesis, we noticed that while control cells grew 1.7 times from S8 to S9 and 1.8 from S9 to S10, mutant cells grew 1.5 and 1.3, respectively (Fig.4G). Thus, it seemed as if the defects in growth observed on the basal surface of mutant FCs increased over time. In fact, when we analysed the basal surface of FCs at later stages, S10A-B, we found mutant cells in which the basal area was hardly visible (Fig.6A-B’). Furthermore, in cross sections of these egg chambers, stained with an antibody Dlg (Fig.6C, C’), or 3D images of wild type and mutant FCs (Fig.6D), it seemed as if, in some cases, wild type cells extended over the basal surface of neighboring mutant ones (arrow in Fig.6C, C’, D). To better understand this phenomenon, we performed live imaging of mosaic S10B egg chambers expressing Resille-GFP and containing *mys*^XG43^ FCs (Fig.6E, E’, Movie 5). We found a striking behavior of the basal surface of wild type cells contacting mutant ones. We observed that, from S9 onwards, the basal surface of control cells that contact mutant ones started to spread anisotropycally over the mutant cells, leading, in some cases, to their disappearance from the focal plane (Movie 5). We did not observe this phenotype in mosaic egg chambers containing control GFP clones (Fig.6F, F’, Movie 6). Anisotropic cell spreading has been proposed to polarize stress fibers ^37^. Accordingly, we found that, while stress fibers are poorly oriented in wild type S10B FCs (Fig.6F, F’), control FCs in contact with *mys*^XG43^ FCs oriented their stress fibers preferentially towards the mutant FCs (Fig.6G, G’). Finally, cell culture experiments have shown that loss of contact with the ECM mediated by integrins results in programmed cell death. However, we found that elimination of integrins in main body FCs did not induce cell death, as tested using an anti-active caspase-3 antibody (Sup. Fig.8). Instead, the integrin mutant FCs displaced by wild type cells are found in the epithelium in a different focal plane, as if they were partially extruded towards the germ line (Fig.6, D).

**Figure 6.**
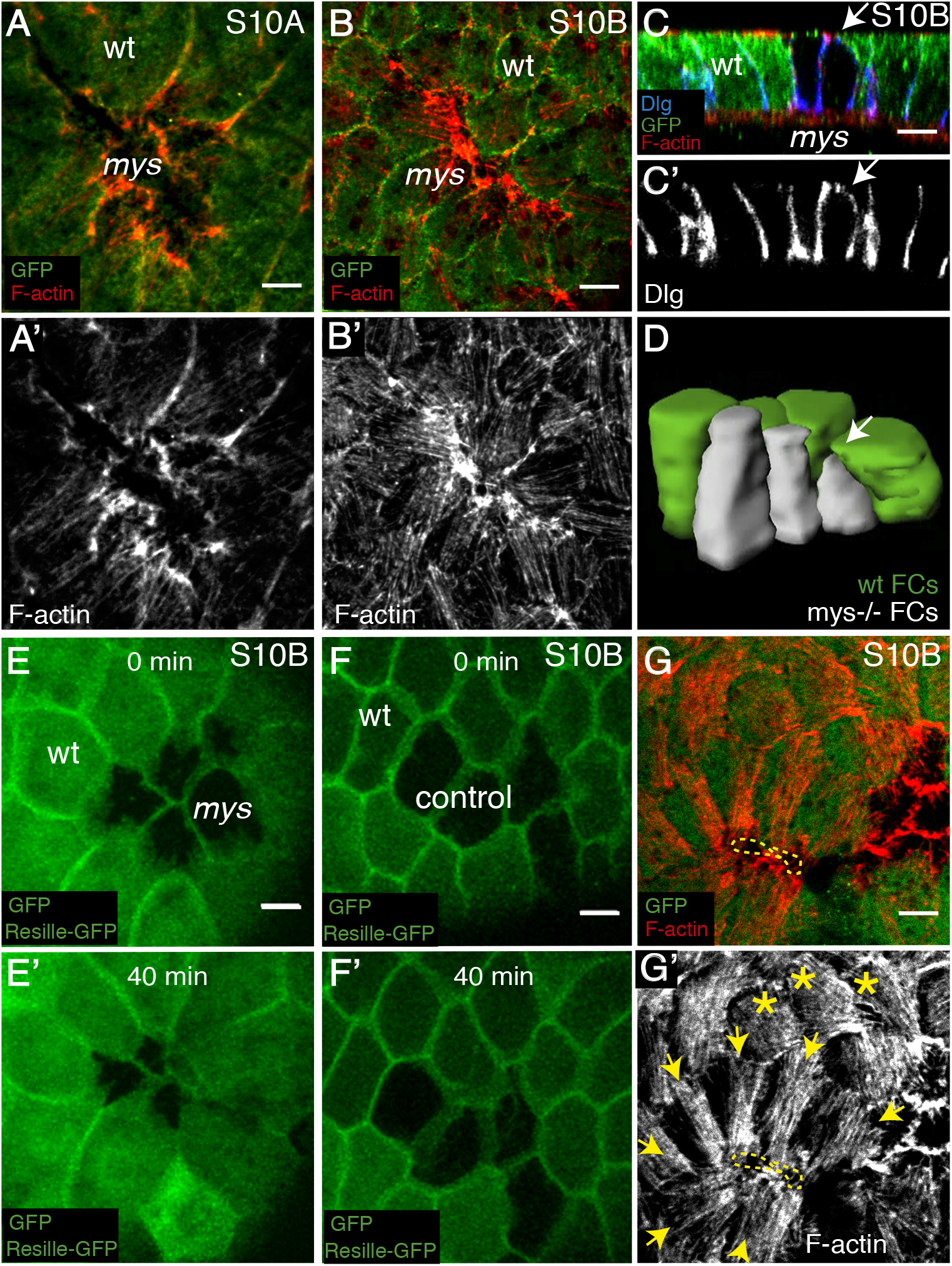
Elimination of integrins in FCs induces spreading of the basal surface of control neighboring FCs. **(A, B)** Basal surface view of S10A **(A, A’)** and S10B **(B, B’)** mosaic follicular epithelia stained with anti-GFP (green) and Rhodamine Phalloidin to detect F-actin (red). **(A, B)** A progressive reduction of the basal surface is observed in *mys* FCs (GFP^-^). **(C)** Lateral view of a S10 mosaic egg chamber stained with anti-GFP (green), Rhodamine Phalloidin (red) and the basolateral polarity marker disc-large (dlg, blue). **(D)** 3D reconstruction of *mys* mutant FCs and surrounding control ones. **(E, F)** Confocal images of live S10B mosaic egg chambers expressing the cell membrane marker Resille-GFP and cytoplasmic GFP and carrying *mys* **(E)** and GFP **(F)** clones. Images were taken with a 40 minutes interval. **(G, G’)** Basal surface view of a S10B mosaic egg chamber carrying a *mys* mutant clone that is almost completely covered by control neighbouring FCs. Yellow asterisks denote control FCs not in contact with mutant cells that orient their fibers randomly (yellow asterisks). In contrast, control FCs enclosing mutant cells orient their basal F-actin fibers towards the mutant ones (yellow arrows). Scale bars, 5 μm.

## Discussion

Actin filaments can assemble into different types of networks that play distinct roles in the cell. Furthermore, morphogenesis seems to be driven by transitions between the different networks. During exit from naïve pluripotency or epithelial-to-mesenchymal transitions actin rearranges from a cortical organization into stress fibers. However, the mechanisms underlying transitions between diverse types of networks are poorly characterized. Equally important to triggering actin network transitions during morphogenesis must be to restrain them in differentiated tissues. Uncontrolled switches in actin networks could result in cell shape and state deregulation, which in turn would affect the morphology and homeostasis of a tissue. Nonetheless, how specific networks are maintained within cells remains elusive. In this work, we have identified the integrins as key regulators of the establishment and maintenance of the different actin networks present in the *Drosophila* FE. Elimination of integrins affects all three types of actin networks found on the basal side of FCs. This has important consequences in both the morphogenesis and the homeostasis of this epithelium.

Stress fibers and focal adhesions are two distinct structures. However, they are highly interdependent. Most of our understating of this relationship comes from studies of cell migration. During cell movement, stress fiber disruption leads to rapid disassembly of the attached focal adhesions and stress fibers diminish when their anchorage sites disassemble ^38^. Likewise, recruitment of actin into stress fibers is impaired when focal adhesion proteins are eliminated ^39^. In addition, actin filament polymerization has been observed at newly formed focal adhesions as the cell moves forward ^40^. Here, we show that elimination of integrins in FCs results in reduced number of basal stress fibers. With time, these stress fibers shorten and collapse. This could be due to the fact that the balance in tension within the stress fibers -as a result of the mechanical resistance of the structures to which they are attached to-is disrupted in integrin mutant cells. This is in agreement with results from cell culture experiments showing that stress fibers contract when separated from their focal adhesions ^41^. In summary, our results suggest that, similar to what happens during cell migration, in static contraction, sites of integrin-mediated adhesion play a key role in the nucleation and maintenance of stress fibers.

Isolated stress fibers retain their contractility ^42^. Likewise, here, we show that actomyosin fibers that are not attached via integrins pulsate. This supports results from cell culture experiments showing localized and stochastic pulses of non-muscle myosin 2 assembly and disassembly in adherent cells, which are independent of integrin-ECM engagement ^32^. Our results also reinforce our recently proposed model, in which periodic basal actomyosin oscillations arise in a cell-autonomous fashion from intrinsic properties of motor assemblies ^16^. However, we find that the spare fibers found in integrin mutant FCs oscillate more stochastically than those of controls. We speculate that in the context of a developing epithelium, cell-ECM interactions mediated by integrins can be required to minimise stochasticity and to guarantee the control of cellular tension, thus allowing the harmonized changes in cell shape required for proper tissue morphogenesis.

We also find here that integrin mutant FCs show higher levels of F-actin in the cortex and in whip-like structures compare to controls. Similarly, detachment of cells from the substratum by trypsinazation causes an increase of F-actin levels in cortical regions ^43^. The mechanisms of redistribution of actin in fibers to a relatively uniform cortex in trypsinized cells remain unclear. The building blocks of stress fibers (actin, myosin and interacting proteins) are constantly exchange with a cytoplasmic pool. Thus, we could propose here that as integrin mutant FCs have fewer stress fibers, there could be an increase in available cytoplasmic actin, which could now transit and associate to the other types of intracellular actin networks. This could also apply to proteins that interact with both F-actin and integrins. For example, moesin, a FERM domain protein that can bind to β1 integrin tails, it is phosphorylated upon cell detachment and it is sufficient to increase cortical F-actin levels and stiffness during mitosis ^43, 44^. Elimination of integrins could result in increased free active Moesin, which in turn would lead to enhanced cortical F-actin. In this scenario, integrins would be required to restrain transitions of actin and/or actin-binding proteins between different networks. Alternatively, integrins could regulate the amount of F-actin and actin interacting proteins present in the different networks independently of each other. Regardless of how they do it, our results show that integrins regulate the amount of F-actin associated to the different types of intracellular networks in the cell, which in turn is key to allow proper epithelia morphogenesis and homeostasis (see below).

Loss of integrin-mediated adhesion results in cell rounding, which has led to suggest that integrins regulate cell surface shape. Treatment of oligodendrocytes with monovalent integrin ligand (RGD-peptide) results in reduced spreading area. This effect is dependent on contractile forces within the cells, as it can be reverted with inhibitors of the ROCK pathway ^45^. This has led to the proposal that integrins regulate cell surface area via a pathway that depends on actomyosin contractility. However, whether this integrin function is involved in the control of cell surface during morphogenesis remains poorly characterized. Here, we show that the basal surface of integrin mutant FCs is smaller than that of controls, which could prevent the correct shaping of the tissue as a whole. In fact, egg chambers carrying integrin mutant FCs do not elongate properly ^12, 17^. Stress fibers are important for the maintenance of shape in crawling cells ^46^. However, more controversial is their role for the overall shape of a non-migrating cell. Our results show that integrin mutant FCs have reduced number of stress fibers and increased cortical F-actin, which is accompanied with a rise in cortical tension. Furthermore, we show that reducing cortical F-actin levels in integrin mutant FCs, without restoring F-actin levels in stress fibers, is sufficient to rescue the reduction in basal surface. This result suggests that, in this context, the main contribution of integrins to the regulation of cell surface area is the modulation of cortical actin contractility rather than the formation and anchorage of stress fibers. This seems to be in contrast with a model from cell culture experiments proposing that cell shape stability depends on the balance between the actomyosin traction forces exerted by the stress fibers and those produced at the adhesion contacts with the ECM ^47^. However, this force balance contributes to cell shape and cytoskeletal stability mainly when cells are cultured on compliant substrates, but not on rigid ones ^41^. In this context, the basement membrane present on the basal side of FCs might be stiff enough to bear the forces produced by the detachment of the stress fibers, upon elimination of integrins, without compromising the overall cellular force balance, and thus, cell shape. Alternatively, cells within tissues are under forces others than those generated by cell-ECM adhesion, such as those mediated by cell-cell adhesion, which might also contribute to balance the reduction in number and detachment of stress fibers caused by elimination of integrins. Finally, our results showing that expression of a dominant negative form of the non-muscle myosin in FCs decreases stress fiber number without reducing cell surface, supports the idea that, in contractile static cells stress fibers might not contribute to cell shape to the same extent they do in migrating cells.

Finally, our results also show that the changes in intracellular actomyosin networks due to the elimination of integrins in a group of cells within a tissue have a non-autonomous effect on the behaviour and cytoskeleton organization of neighbouring wild type cells. This effect is similar to that observed upon tissue wounding. During wound healing, cells around the wound sense the damage and form actin-based cellular protrusions that allow them to crawl forward, repair the wound and maintain tissue homeostasis. Likewise, here, we find that control cells surrounding integrin mutant FCs spread their basal surface anisotropically over that of mutant cells. Furthermore, with time, the basal side of control FCs ends up covering the space initially occupied by that of mutant cells. During this asymmetric cell spreading, control FCs re-orient their stress fibers towards the mutant ones, supporting a model from cell culture experiments pointing to asymmetric cell spreading as a mechanism to regulate stress fibers polarity ^37^. Data from cell culture experiments have also led to suggest that availability of free space is sufficient to trigger cell migration in the absence of mechanical injury ^48^. Thus, as integrin mutant FCs most likely detach from the basement membrane, we would like to propose that, similar to the way epithelial cells sense a breach in a tissue and act to repair it, they could sense available free ECM space and cover it. In doing so, wild type cells pushed integrin mutant ones inwards. Loss of α2β1 integrin expression results in increased extravasation in breast and prostate cancer ^49^. The phenomenon here described could be a mechanism to facilitate the invasion and metastasis of tumor cells with low levels of integrins.

Intracellular actin networks can organize in diverse patterns. In addition, although they normally localize to precise regions of the cells, they are rarely independent and often their dynamics influence each other. In fact, the reorganization of a given structure can promote the formation of another, conversions that govern many morphogenetic processes. In addition, a wide range of diseases, including neurological and musculoskeletal disorders as well as cancer, result from defects in actin networks assembly and uncontrolled transitions. Here, we propose that cell-ECM interactions mediated by integrins control epithelia morphogenesis and homeostasis by sustaining the different types of intracellular actin networks. Identifying new regulators either to restrain or to trigger transitions between different types of actin networks is crucial to fully comprehend morphogenesis and the cellular and molecular basis of some pathologies.

## Methods

### Drosophila stocks and genetics

The following fly stocks were used: *mys^11^* (also known as *mys^XG43^* ^23^, Sqh-GFP ^50^, Sqh-mCherry ^24, 50^ from Bloomington *Drosophila* Stock Centre, UAS-*abi*RNAi (DGRC-Kyoto 9749R), the follicle stem cell driver *traffic jam*-Gal4 (*tj-gal4*, ^51^) and the cell membrane marker Resille-GFP ^52^. The protein trap line mys-GFP is a gift from Prof. N. Brown. The *e22c-gal4* driver is expressed in the follicle stem cells in the germarium and was therefore combined with *UAS-flp* to generate *mys* FC clones. To visualise cell membranes in *mys* mutant clones, *mysXG43FRT101*/FMZ; Resille-GFP females were crossed to *ubiquitin*-GFPFRT101; *e22c-gal4 UAS-flp/CyO* males. To analyse myosin dynamics in *mys* mutant clones, *mysXG43FRT19A/FMZ;* Sqh-GFP or *mysXG43FRT101/FMZ;* ResilleGFP:Sqh-mCherry/CyO females were crossed to *nlsRFP FRT19A; e22c-gal4 UAS-flp/CyO* and ubiquitin-GFPFRT101; *e22c-gal4 UAS-flp/CyO* males, respectively. To study F-actin distribution and dynamics an ubiquitin-lifeactinYFP construct (described below) was generated and recombined with Resille-GFP. To analyse actin dynamics in *mys* mutant clones, *mysXG43FRT101*/FMZ; *ubiquitin*-lifeactinYFP:Resille-GFP/CyO females were crossed to *ubiquitin*-GFPFRT101; *e22c-gal4 UAS-flp/CyO* males. To analyse the behaviour of GFP control clones, we crossed FRT101/FMZ; Resille-GFP females to *ubiquitin*-GFPFRT101; *e22c-gal4 UAS-flp/CyO* males. For the rescue experiments, we used the heat shock flipase (*hs-flp*) system ^53^ to generate follicle cell mutant clones and *tj*-Gal4, recombined with Resille-GFP to visualize cell membranes, to express the UAS-*abi*RNAi. *mysXG43FRT101/FMZ;* UAS-*abi*RNAi/FMZ females were crossed to *hs-flpGFP FRT101*; *tj*-Gal4:Resille-GFP males. The heat shock was performed at 37°C for 2 h during third instar larvae and newly hatched females. Flies were kept at 25°C and yeasted for 2 days prior to ovary dissection.

### Ubiquitin-lifeactinYFP construct

Sequences for the actin-binding peptide lifeactin tagged with the fluorescent protein YFP with optimized codon use for *Drosophila* and KpnI and NotI ends were designed. They were synthetized *“in vitro*” by Sigma. The sequences were cloned into the polylinker of the pWR-pUbq transformation vector. This vector contains a poliubiquitin promoter and a selectable marker *mini-white*+. The plasmid was introduced into the germ line of *w*(1118) flies by standard methods by the company BestGene Inc. and several independent transgenic lines were isolated.

### Immunohistochemistry

Flies were grown at 25 °C and yeasted for 2 days before dissection. Ovaries were dissected from adult females at room temperature in Schneider’s medium (Sigma Aldrich). The muscle sheath that surround ovarioles were removed at this moment. After that, fixation was performed incubating egg chambers for 20 min with 4% paraformaldehyde in PBS(ChemCruz). Samples were permeabilized using PBT (phosphate-buffered saline+1% TritonX100). For actin labelling, fixed ovaries were incubated with Rhodamine-Phalloidin (Molecular Probes, 1:40) for 40 min. The following primary antibodies were used: chicken anti-GFP (1/500, Abcam) and mouse anti-Dlg (1/50, DHSB, Iowa). Fluorescence-conjugated antibodies used were Alexa Fluor 488 and Alex Fluor 647 (Life Technologies). Samples were mounted in Vectashield (Vector Laboratories) and imaged on a Leica SP5 MP-AOBS.

### Time-lapse image acquisition

For live imaging 1–2 days old females were fattened on yeast for 48–96 hours before dissection. Culture conditions and time-lapse microscopy were performed as described in ^54^. Ovarioles were isolated from ovaries dissected in supplemented Schneider medium (GIBCO-BRL). Movies were acquired on a Leica SP5 MP-AOBS confocal microscope equipped with a 40 × 1, 3 PL APO oil and HCX PL APO lambda blue 63x 1.4 oil objectives and Leica hybrid detectors (standard mode). Z-stacks with 20-23 slices (0.42 μm interval) were taken to capture the entire basal surface of the cells with time points every 30 seconds up to 1 hour.

### Laser ablation

Laser ablation experiments were performed in an inverted Axiovert 200 M, Zeiss microscope equipped with a water-immersion lens (C-Apochromat 633 NA 1.2, Zeiss), a high-speed spinning-disc confocal system (CSU10, Yokogawa), a cooled B/W CCD digital camera (ORCA-ER, Hamamatsu) and a 355 nm pulsed, third-harmonic, solid-state UV laser (PowerChip, JDS Uniphase) with a pulse energy of 20 mJ and 400msec pulse duration. A Melles Griot ArIon Laser (l = 488 nm, 100 mW) was used for excitation of enhanced green fluorescent protein. To analyze the vertex displacements of ablated cell bonds, we first averaged the vertex distance increase from different ablation experiments (L) using as L_0_ the average of distance of the vertex ten seconds before ablation. Changes in vertex distance before ablation are caused by the movement of cell membranes. Images were taken before and after laser pulse every 0,8 seconds for a period of 10 seconds. The initial velocity was estimated as the velocity at the first time point (t1 = 0,8 s). Standard errors were determined.

### Image processing and data analysis

For quantification of basal myosin and actin dynamics over time, maximal projections of confocal stacks were produced to cover for egg chamber curvature. Integrated intensity of myosin and actin were quantified for manually selected regions using ImageJ software. The background value taken from cell-free regions was subtracted from all data series. Data were subjected to Gaussian smoothening with s=3, σ=3. The distribution of oscillation periods was obtained by measuring the intervals between each pair of two adjacent peaks. Actin and myosin intensity changes in both wild type and *mys* mutant cells were obtained by averaging the difference between the maximum and the minimum fluorescence intensity for each oscillation. To calculate the period of myosin and actin oscillation, a Matlab script described in ^16^ was used to measure the power spectrum density of the signal using one-dimensional Fourier transform of the autocorrelation function.

Number of stress fibers was calculated using ImageJ software. First, the entire basal surface of the cell, excluding the cortical region, was outlined. Then, a line extending across this basal surface, on its central region, perpendicular to the actin bundles was drawn. “Plot profile” tool was employed to quantify the fluorescence intensity of the peaks along this line. To quantify peaks in mutant FCs, only peaks greater than one value of SD below the mean intensity of those found in wildtype cells were considered. Number of peaks within a cell divided by the cell area was used to compute peak density (Number of picks/μm). Measurement of whole fluorescence intensity was done by dividing the mean of all included pixels intensity by the outlined cell area. Since we needed to adjust laser intensity in each sample to properly visualize actin bundles, due to staining heterogeneity, the ratio between the fluorescent intensities of *mys* and wild type cells was plotted.

To determine cortical actin intensity across cell edges, custom 2-micrometers-long scripts in Image J were used. Most representative curves of actin fluorescence intensity at each point along the script were used to show the magnitude and distribution of actin at the cell edges. The whole fluorescence intensity along the script was normalized with respect to wild type values.

Cell area data were calculated using Imaris (Bitplane). The whole basal surface of the cell was outlined using Resille-GFP as cell membrane marker.

## Acknowledgements

We thank the Bloomington and Kyoto Stock Centres for fly stocks and reagents. We are grateful to A. Gonzalez-Reyes for useful comments on the manuscript. Research in our laboratories is funded by the Spanish Ministerio de Economía y Competitividad and the FEDER programme (BFU2013-48988-C2-1-P and BFU2016-80797-R) to M.D. M-B.) and by the Junta de Andalucía (Proyecto de Excelencia P09-CVI-5058). C. S-C M and A. V-E were supported by FPU and FPI Fellowships, respectively (Ministerio Español de Economía y Competitividad). IMP was supported by the BBSRC, the University of Cambridge and Queen Mary University of London

## Author contributions

C. S-C M and A. V-E performed experiments, analysed data and contributed to the writing of the manuscript. D.G.M. generated the Fourire transform graphics and make helpful comments on the manuscript. I.P. suggested ideas and contributed to the writing of the manuscript. M. D. M.-B. provided financial support, designed research, analysed data and wrote the manuscript.

