## Supplementary figures and images for "Integrins control tissue morphogenesis and homeostasis by sustaining the different types of intracellular actin networks"

### Supplemental Fig.1

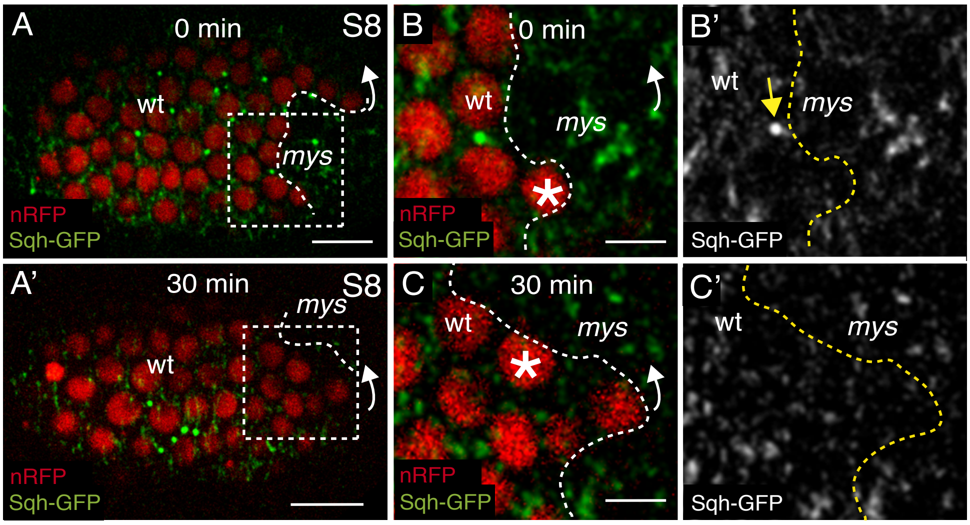


Suplemmentary Figure 1

### Supplemental Fig.2

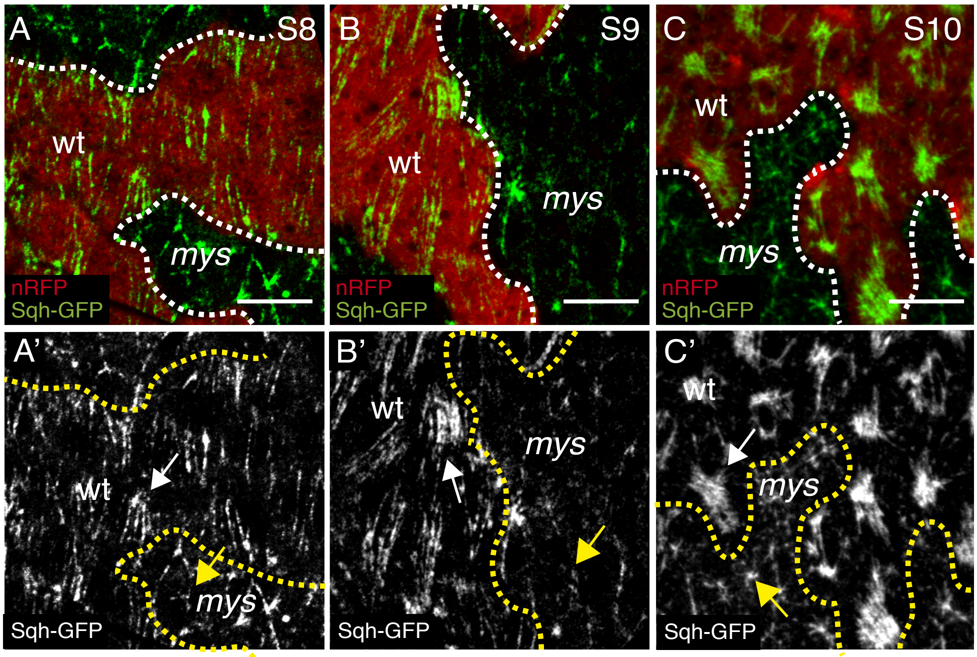


Supplementary Figure 2

### Supplemental Fig.3

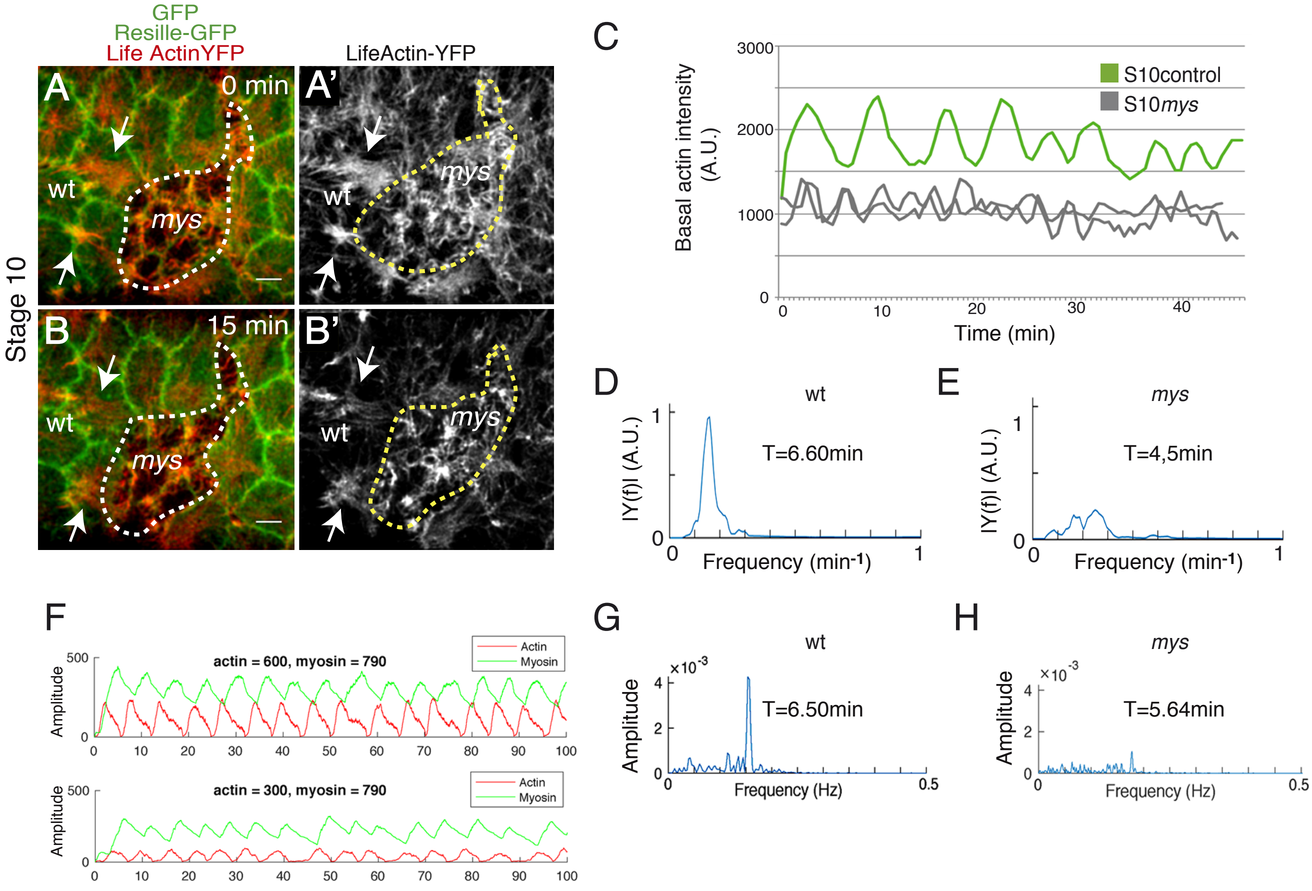


Supplementary Figure 3

### Supplemental Fig.4

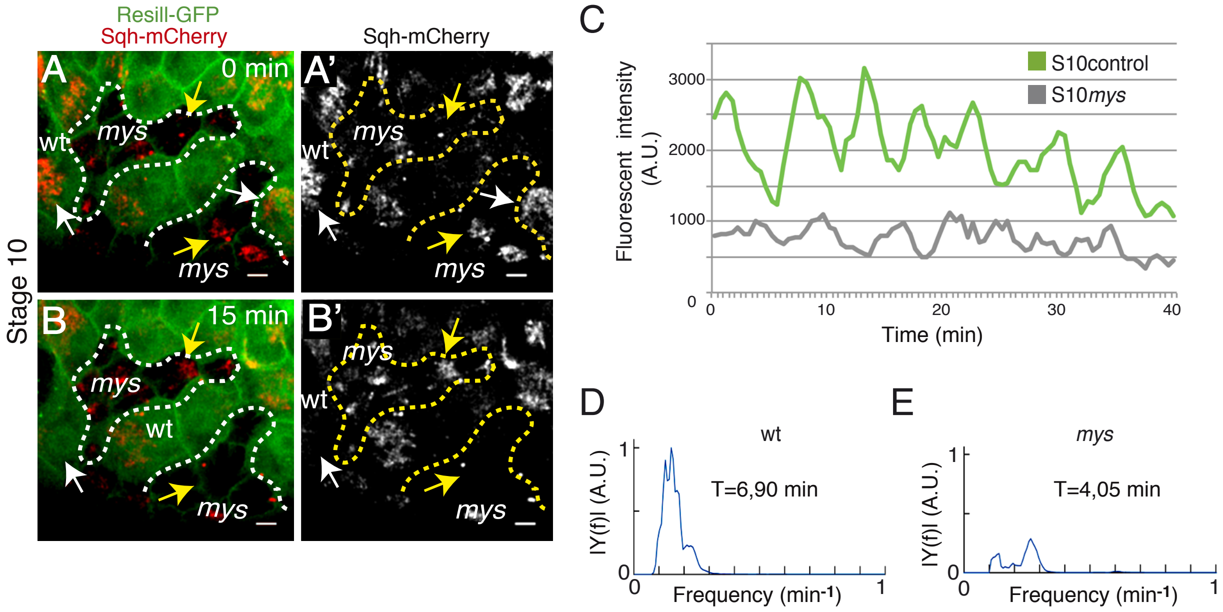


Supplementary Figure 4

### Supplemental Fig.5

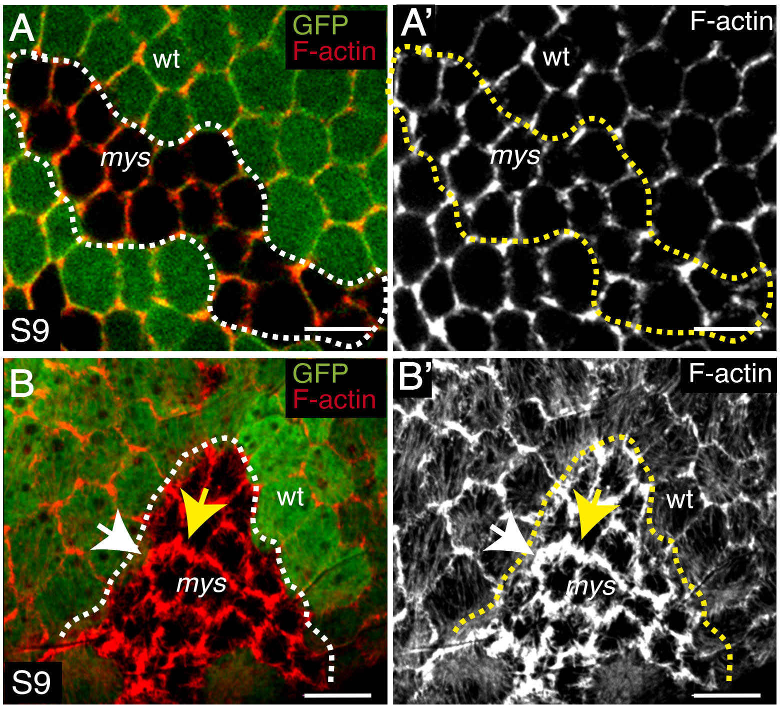


Supplementary Figure 5

### Supplemental Fig.6

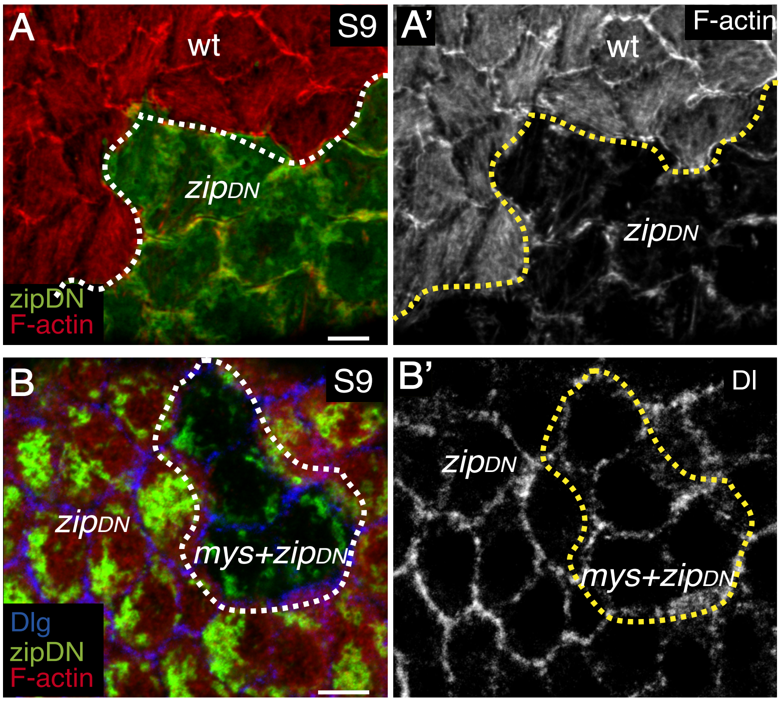


Supplementary Figure 6

### Supplemental Fig.7

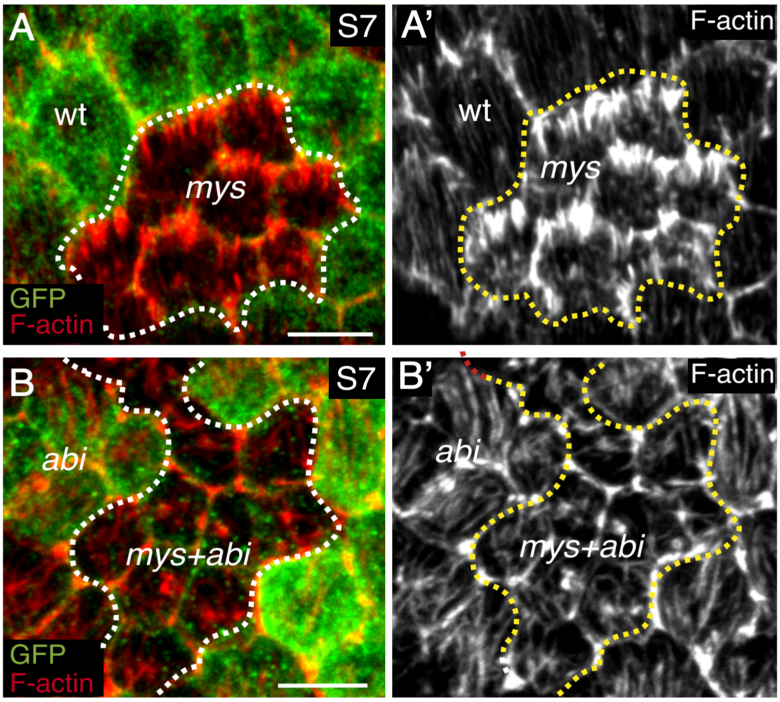


Supplementary Figure 7

### Supplemental Fig.8

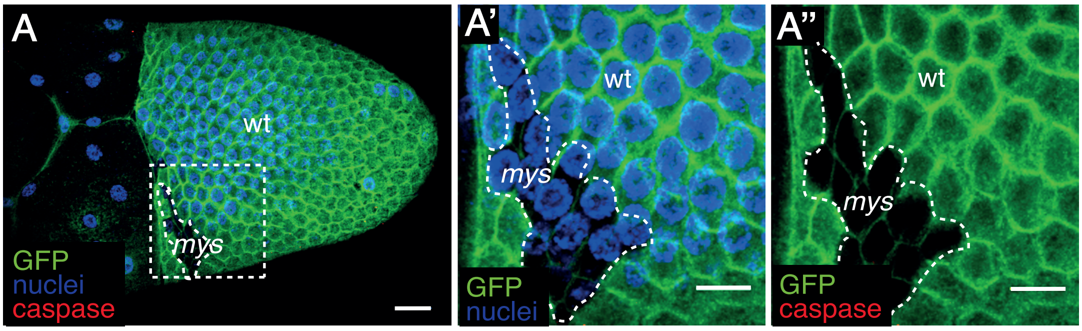


Supplementary Figure 8
